# Quantifying floral rewards: Comparative evaluation of nectar extraction techniques

**DOI:** 10.64898/2026.11.16.766450

**Authors:** Vidisha Bansal, Kenneth Kuba, Gaya M. ten Kate, Carmen A. Nebauer, Paula Prucker, Fabian A. Ruedenauer, Ling Xu, Ada D. Özkir, Sara D. Leonhardt

## Abstract

1.) Nectar is a crucial resource for a wide range of insect species and one of the driving forces shaping plant-pollinator interactions. Flowers have evolved diverse strategies to restrict nectar access, ensuring only beneficial flower visits resulting in successful pollination. This range of adaptations, often morphological, makes it difficult for researchers to collect nectar in a standardised manner. In this study, we compare common methods used for nectar extraction, with a focus on applicability across several plant families

2.) We assessed pre-treatment methods widely used before nectar extraction, such as pollinator exclusion by bagging flowers, storing flowers overnight in water, and standing nectar crop (unmanipulated control). Moreover, we compared the effectiveness of two commonly used nectar extraction techniques, i.e., centrifugation and microcapillaries, across a range of flower morphologies. We compared nectar volume, sugar concentration, and total sugar content obtained by different methods for 23 flower species from 16 plant families.

3) We show that pollinator exclusion by bagging is an effective measure to obtain maximum nectar volume regardless of plant growth type. The nectar sugar concentration of herbaceous and woody species showed pre-treatment specific variation. Extraction with a centrifuge consistently yielded more nectar volume without altering the sugar concentration, while microcapillary extraction was not effective for most flower morphologies, especially lip (e.g. *Salvia pratensis*) and disc-shaped flowers (e.g. *Rosa canina*).

4.) Our results provide a foundation for the standardisation of floral nectar extraction protocols, improving comparability across studies, and facilitating broader ecological and evolutionary research that depends on consistent and reliable nectar sampling.

## 1. Introduction

Nectar is a carbohydrate-rich aqueous secretion produced primarily by flowers and important for interaction outcomes between plants and pollinators (Heil, 2011; Parachnowitsch et al., 2019; Potts et al., 2004). Nectar volume and sugar concentration determine the energy provided to pollinators foraging on plants and can strongly influence visitation and thus pollination effectiveness (Fornoff et al., 2017; Heinrich, 2004; Potts et al., 2004). Information on nectar availability across different flowering plant species is therefore crucial for understanding resource availability and quality in the foraging landscapes of pollinators (Parreño et al., 2022), which is a prerequisite for combatting the ongoing pollinator decline and for informing conservation and restoration efforts (Goulson et al., 2015).

Nectar is produced from photosynthetically derived sugars and secreted by specialised glands, called nectaries, that are located either within the flower structures (i.e., floral) or on other parts of the plant (i.e., extrafloral) (Marazzi et al., 2013; Pacini et al., 2003). Nectar composition and volume varies widely among plant species and is influenced by environmental, physiological and evolutionary factors (Biella et al., 2022; Descamps et al., 2021; Leonhardt et al., 2024; Parachnowitsch et al., 2019; Venjakob et al., 2022). Weather conditions, i.e., temperature, solar radiation and water availability additionally affect nectar secretion dynamics and evaporation rates (Descamps et al., 2021; Heil, 2011; Kram & Carter, 2009; Plos et al., 2023), often resulting in fluctuations in nectar production throughout the day (Bloch et al., 2017; Silva et al., 2004; Witt et al., 1999). Moreover, plant growth type, i.e., herbaceous or woody, can influence nectar availability, as some woody species have carbon-autonomous branchlets that support fruit and flower growth even if the connection to the roots is severed (Hoch, 2005).

The observed strong variation in nectar production between and within plant species calls for efforts to standardise measurements, which has led to the adoption of a few commonly used flower treatments prior to nectar extraction. A common approach to obtain measurements of total nectar quantity is, for example, bagging flowers (i.e., covering them with a nylon mesh bag) to exclude pollinators and allow nectar to accumulate (Corbet, 2003; Pioltelli et al., 2024; Venjakob et al., 2022; Wyatt et al., 1992). Similarly, cutting plants and storing them in water overnight, out of reach of pollinators, is another pre-treatment method (see, for example, Bansal et al., 2026). Alternatively, scientists measure the standing crop, i.e., extract nectar in situ from undisturbed flowers, which is considered the ecologically relevant availability of nectar at a certain point in time (Corbet, 2003). The effects of each of these methods on nectar quality and quantity, and thus the risk of introducing methodological bias, are largely unknown. Likewise, the effects of various techniques for nectar extraction e.g., capillaries, wicks, filter papers, or centrifuges, with potential tool-specific biases on nectar measurements remains unexplored (Corbet, 2003; Pioltelli et al., 2024). Using capillaries requires the nectar reservoir to exceed a minimum threshold to collect pure nectar, and extreme caution must be exercised to avoid damaging surrounding plant tissue and plant fluids that dilute the nectar (Corbet, 2003). However, many plant species, e.g., *Bellis perennis,*exhibit structures that are too thin or cover nectaries, rendering nectar inaccessible to most capillaries (Corbet, 2003). To overcome these challenges, some studies have reported the use of handcrafted ultra-thin capillaries for nectar extraction (Corbet, 2003). Alternatively, flowers can be centrifuged to extract nectar (Bansal et al., 2026; Na et al., 2025; Pioltelli et al., 2024; Swanson & Shuel, 1950). A systematic comparison between using capillaries and a centrifuge, however, is still missing, preventing the development of a standardised, comparable, and repeatable approach to nectar extraction across diverse flower types (Michelot-Antalik et al., 2025).

Floral morphology, i.e., the shape and form of floral structure, determines the location and shape of nectaries and nectar-containing compartments (Keasar & Bodner, 2025). Two prominent factors that influence flower morphology are pollination syndromes and phylogeny (Bailes et al., 2023; Caruso, 2000; Chartier et al., 2014; Gervasi & Schiestl, 2017; Kugler, 1970; Newman et al., 2014). While phylogeny explains morphological similarities in closely related flowers, pollination syndromes explain similarities within flowers that are visited/pollinated by similar groups of animals (Fenster et al., 2004; Kugler, 1970). For example, stalked disc flowers are often associated with moths (Lepidoptera) and Bombyliidae (Diptera), both of which collect nectar with long-tubed probosces (Krenn et al., 2005; Kugler, 1970). On the other hand, *Rosa canina* has an open disc flower (Kugler, 1970) that is accessible to most insects, including short-tongued ones. Thus, different floral shapes constrain nectar accessibility differently and thereby affect extraction efficiency, underscoring the need to account for floral shape in methodological comparisons.

Despite widespread use and some previous comparisons of different nectar extraction techniques (Michelot-Antalik et al., 2025; Pioltelli et al., 2024), the effect of pre-treatment methods in combination with extraction tools, plant growth type, and flower morphology remains unknown. In our study, we therefore compared the most common pre-treatment methods for nectar extraction across 23 species to determine if nectar extraction using different approaches is comprehensive and consistent. To provide a foundation for a standardised protocol for nectar extraction, we conducted standardised flower collection using three defined pre-treatments, i.e., overnight water storage, pollinator exclusion by bagging, and standing nectar crop (unmanipulated control), and extracted nectar via two methods: microcapillaries and centrifugation. We tested differences among these methods in measured nectar volume, sugar concentration, and sugar content. We further explored the effects of plant growth type and flower shape on nectar extraction by different techniques, and the influence of phylogeny on nectar quantity and quality. Informed by the results we provide a step-by-step guide for nectar extraction from various flowers using centrifugation and a 3-D printable design of mesh sieves for preventing contamination in the nectar sample.

We hypothesised that different pre-treatments would affect the nectar volume. Although bagging flowers prevents nectar consumption by pollinators, it does not standardise the extent of water loss from nectar due to evaporation. Plucking plants at different times and keeping them in water overnight should, in turn, increase the overall nectar volume and thus reduce the sugar concentration. Regarding growth type, we expected woody plants to be less affected by pre-treatments and exhibit lower variability than herbaceous species, as they have a pronounced secondary vascular system and storage capacity within individual branchlets, thereby aiding the mobilisation of fresh nectar (Hoch, 2005).

We further hypothesize that nectar extraction with centrifugation yields high nectar volume, while nectar sugar concentrations are not affected. However, as the quantity of nectar extracted per flower depends largely on flower morphology, we expected microcapillary extraction of nectar in lip, and stalk disc flowers would yield results similar to those of the centrifugation method. Additionally, we expected the centrifugation method to yield more nectar in plants with composite, and disc-shaped flowers.

## 2. Materials and methods

### 2.1 Plant collection

Plants were collected on Weihenstephan campus of the Technical University of Munich (TUM) in Freising, Germany. Sampling took place from April to June 2025. The selection of plant species was based on availability, taxonomic diversity, and flower morphology, including both herbaceous (annual or perennial plants) and woody species (shrubs and trees). Plants were identified to species level using a field guide for the identification of flowering plants (Aichele & Golte-Bechtle, 1980) or Flora Incognita (Mäder et al., 2021). Each species was collected locally where it flowered abundantly. Flowers from six plant individuals per species and pre-treatment were measured to account for intraspecific variation. Flower collection took place during dry weather, and rainy days were avoided as precipitation affects the nectar production in plants (Carroll et al., 2001; Descamps et al., 2018; Glenny et al., 2018), resulting in diluted nectar with low sugar content (Enkegaard et al., 2016). Most species were collected in the morning between 8 a.m. and 12 p.m. except for *Berberis verruculosa* which was collected at 4 p.m. Flowers collected for all pre-treatments were plucked below the second node and transported for further processing in water.

### 2.2 Flower morphology categorisation

All plant species were assigned to flower morphology categories based on Kugler, (1970) using information on flower characteristics according to the Bundesamt für Naturschutz via the FloraWeb database (Bundesamt für Naturschutz (BfN), 2026; Klotz et al., 2002). If a species had no database entry, information was inferred from a closely related species with an existing entry, visual inspection and comparison with the original literature (Kugler, 1970). Floral morphology is defined in detail by Kugler, where each category is divided into multiple subcategories (e.g. disc flowers with nectar ± hidden nectaries at base of stamens). We omitted the subcategories and used only the main category (e.g. disc and bowl-shaped flowers) to create statistically testable groups. Consequently, all species were assigned into five distinct floral morphologies (Table 1): disc and bowl-shaped flowers (Fig. 1a), stalk disc flowers (Fig. 1b), lip flowers (Fig. 1c), papilionaceous flowers (Fig. 1d) and composite flowers (Fig. 1e). Although *Trifolium pratense* has papilionaceous florets (Fig. 1e), it is classified as a composite flower for this study as the nectar traits were extracted at the inflorescence level rather than per individual floret.

**Figure 1:**
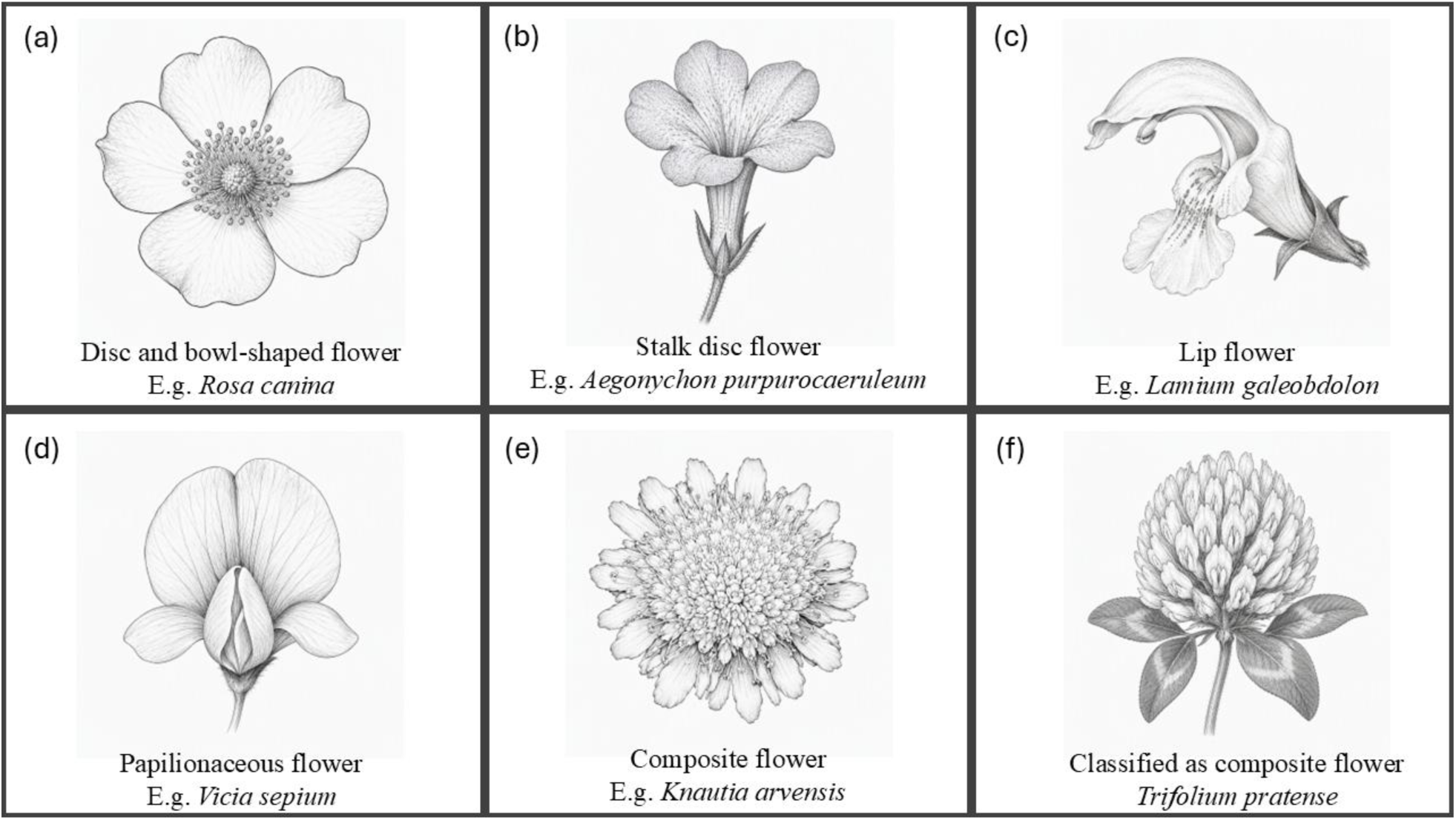
Flower morphology categories following Kugler (1970). All species were classified according to this scheme, except *Trifolium pratense*, which was categorised as a composite flower rather than a papilionaceous flower because nectar traits were measured at the inflorescence level. Image generated using Microsoft Copilot, 2026 (https://copilot.microsoft.com/) based on photographs clicked by the authors.

**Table 1:** Details about the plant species sampled, their taxonomic family, growth type, the flower morphology as well as the centrifugation speed (rpm) and the duration of centrifugation (s) needed to extract the maximum nectar quantity without damaging the flower. Species indicated with an asterisk (*) did not have their floral morphology listed on FloraWeb database (Klotz et al., 2002; floraweb.de). Their floral morphology was examined and assigned to the categories described in Kugler, (1970) by the authors.

| Species | Family | Flower morphology | Growth type | Centrifugation speed (rpm) | Centrifugation duration (s) |
|---|---|---|---|---|---|
| *Aegonychon purpurocaeruleum* | Boraginaceae | Stalk disc | Herbaceous | 10000 | 10 |
| *Aesculus hippocastanum* | Sapindaceae | Lip | Woody | 8000 | 30 |
| *Allium ursinum* | Amaryllidaceae | Disc flower | Herbaceous | 8000 | 20 |
| *Berberis verruculosa* | Berberidaceae | Disc flower | Woody | 10000 | 20 |
| *Bunias orientalis* | Brassicaceae | Disc flower | Herbaceous | 4000 | 10 |
| *Colutea orientalis* | Fabaceae | Papilionaceous | Woody | 10000 | 20 |
| *Cornus sanguinea* | Cornaceae | Disc flower | Woody | 8000 | 10 |
| *Deutzia crenata\** | Hydrangeaceae | Disc flower | Woody | 8000 | 30 |
| *Dianthus carthusianorum* | Caryophyllaceae | Stalk disc | Herbaceous | 8000 | 10 |
| *Geranium pyrenaicum* | Geraniaceae | Disc flower | Herbaceous | 6000 | 10 |
| *Knautia arvensis* | Caprifoliaceae | Composite | Herbaceous | 6000 | 10 |
| *Kolkwitzia amabilis\** | Caprifoliaceae | Stalk disc | Woody | 10000 | 20 |
| *Lamium galeobdolon* | Lamiaceae | Lip | Herbaceous | 10000 | 20 |
| *Lotus corniculatus* | Fabaceae | Papilionaceous | Herbaceous | 8000 | 10 |
| *Onobrychis viciifolia* | Fabaceae | Papilionaceous | Herbaceous | 10000 | 10 |
| *Rosa canina* | Rosaceae | Disc flower | Woody | 8000 | 30 |
| *Salvia pratensis* | Lamiaceae | Lip | Herbaceous | 10000 | 10 |
| *Styrax japonicus* | Styracaceae | Disc flower | Woody | 10000 | 20 |
| *Trifolium pratense* | Fabaceae | Composite | Herbaceous | 6000 | 10 |
| *Veronica chamaedrys* | Fabaceae | Papilionaceous | Herbaceous | 10000 | 10 |
| *Vicia sepium* | Plantaginaceae | Lip | Herbaceous | 8000 | 70 |
| *Vicia villosa* | Fabaceae | Papilionaceous | Herbaceous | 8000 | 60 |
| *Weigela florida* | Caprifoliaceae | Stalk disc | Woody | 12000 | 30 |

### 2.3 Pre-treatments

Six plant individuals were selected in a random manner for each pre-treatment method. Replicates refer to either one flower (e.g., *Rosa canina*), several florets (e.g., *Vicia villosa*), or a whole composite flower (e.g., *Trifolium pratense*), depending on the species. Comparable number of plant species with woody and herbaceous growth type with different flower morphologies were sampled for the study. All nectar trait measurements were calculated per floret or composite flowerhead that were recorded during sampling. The following three pre-treatments were established:

#### Control (Standing crop)

Untreated flowers were harvested on the day of nectar extraction. They were immediately transferred to the lab within 10 – 15 minutes for extraction without further manipulation.

#### Bagging

Plant sites were visited two to five days before nectar extraction, and flower buds or flower stalks/composites with plenty of unopened florets were bagged with a fine nylon mesh bag (8 cm x 6 cm, mesh size 0.38 mm) to exclude pollinators and let nectar accumulate in the flowers (Fig. 2a).

**Figure 2:**
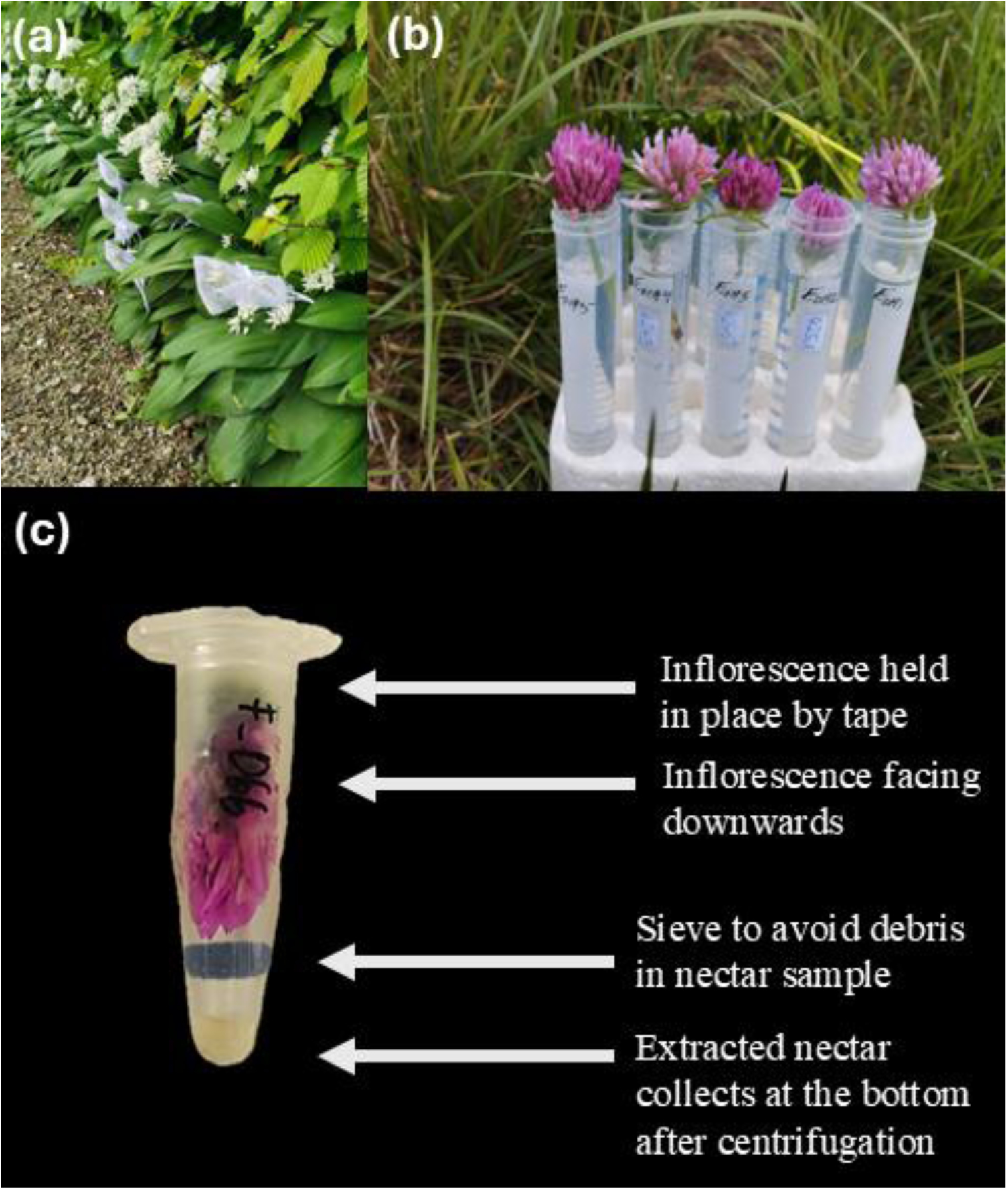
Two methods were used as pre-treatment before nectar extraction and compared with control (standing crop): nectar exclusion by bagging (a) and overnight water storage (b). The flowers were taped and held in place upside-down in Eppendorf or Falcon tubes and a sieve was used to avoid debris in the extracted nectar samples (c).

#### Overnight water storage

Plants were collected the day before the nectar extraction. The collected flowers were kept overnight in a well-lit room at 22°C in tap water to allow nectar production and accumulation (Fig. 2b). Nectar was extracted the following morning.

### 2.4 Extraction

Two extraction methods were tested in this study:

#### Capillary

Before extraction, nectaries were located based on pollinator observations and dissection of flowers. Capillaries were selected based on the corolla tube length of the florets. Small glass capillaries (Hirschmann minicaps® 2 µl, 5 µl, and 10 µl) were used for disc flowers (e.g. *Rosa canina*), and long glass capillaries (CAMAG® 5 µl) were used for flowers with long corolla tube length (e.g. *Aegonychon purpurocaeruleum*). Flowers were sampled using microcapillaries until the entire nectar volume had been extracted.

#### Centrifuge

A adaptive centrifuge extraction of floral nectar was used, a detailed description is provided with the resulting step by step guide https://doi.org/10.5281/zenodo.24201322. For centrifugation (Eppendorf tubes: Centrifuge 5430, Eppendorf AG, Hamburg, Germany; Falcon tubes: Sigma 6-16S, Sigma Laborzentrifugen GmbH, Osterode am Harz, Germany), flowers were placed in Eppendorf (1.5 ml) or Falcon tubes (15 ml), depending on the flower diameter. Clear tape was used to wrap florets together at their base, thus reducing the overall size. All flowers were inserted into the tubes facing towards the bottom of the tube and the tape at the top was fixed in place using the Eppendorf tube lid (Fig. 2c). Additionally, we added sieves to the tubes to reduce nectar contamination with debris https://doi.org/10.5281/zenodo.22875506. We adapted the approach proposed by Pioltelli et al. (2024) by designing PLA 3-D printed sieves in three sizes that fit exactly into common centrifuge tubes (see data availability statement). These sieves were designed to sit above the tips of the tubes and with mesh size ranging between 0.4 - 0.5 mm (Fig. 2c).

First, we established the centrifugation parameters, i.e., speed and time, chosen with the slowest ramp (rotational amplitude), acceleration and deceleration time. The flowers were centrifuged in a succession of runs, and the effects on the flowers were visually evaluated as follows:

#### Speed

Three tubes with flowers were centrifuged at 2000 rpm for 10 seconds. The tubes were inspected for (i) signs of nectar, e.g., droplets of transparent liquid or smears along the tube wall, and (ii) flower state, if the structure is still intact. If the flowers were intact, the speed was increased by 2000 rpm, and inspection was repeated. This procedure was continued until the flowers showed signs of damage (e.g., ruptures). If the flowers were damaged, the last recorded speed with the flower in good condition was subsequently used to extract nectar from other replicates for the final dataset.

#### Time

The centrifuge was reloaded with three tubes of fresh flowers. We used the previously determined speed and centrifuged the flowers multiple times for 10 seconds. While ensuring the flowers remained intact, we checked for nectar droplets after every 10 s interval. If no increase in nectar amount was observed, the trials were stopped, and the combined time of all the intervals was chosen as the extraction time. In cases where flowers were damaged or loosened between intervals, the lower time limit was used for other replicates. This method was adjusted for all plant species (Table 1). After centrifugation, flowers were carefully removed, and tubes were centrifuged again at 10,000 rpm for 10 seconds to concentrate all nectar at the base of the tubes for easy collection via glass microcapillaries.

### 2.5 Nectar trait measurements

The nectar was collected either directly using glass microcapillaries or by collecting the nectar extracted at the bottom of the Eppendorf or Falcon tubes after centrifugation. To obtain the nectar volume per flower, the following formula was used:

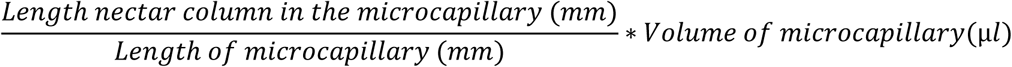

The length of the microcapillary and the nectar column in the microcapillary were measured using a digital calliper (Burg-Wächter Precise Digital Vernier calliper - PS 7215, Wetter, Germany). At least 5µl of nectar was transferred to a refractometer (Eclipse Professional, 0-50 °Brix and 45-80 °Brix, Bellingham + Stanley, Tunbridge Wells, UK) using pipette bulbs (Hirschmann™ Pipette Bulbs) to measure the relative amount of sucrose solution (°Brix) for each sample. As a minimum of 5µl was needed for a reliable measurement of sugar concentration, nectar was pooled across several replicates from the same plant species to obtain a sugar concentration measurement. Sugar mass was then calculated using the following formula:

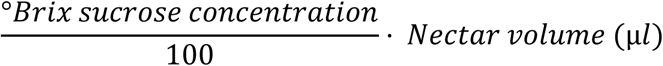

### 2.6 Statistical analysis

The entire data analysis was carried out in R version 4.2.2 (R Core Team, 2025). Generalised linear mixed-effects models (GLMMs) based on the “lme4” package (Bates et al., 2015) were used to assess the effect of pre-treatments and their interaction with growth type, and the effect of extraction methods and their interaction with flower morphology on nectar volume, sugar concentration, and sugar mass for each plant species. The temperature and the weather at the time of collection, as well as plant species were included as random effects. The distribution of the residuals and the homoscedasticity of the models were determined using the “DHARMa” package (Hartig et al., 2024). The explanatory variables were log-transformed to improve model fit. As we were checking for interactive effects in our models, type III Wald’s tests were conducted to determine the significance of explanatory variables on the response variables using the “car” package (Fox & Weisberg, 2019). Post-hoc tests were conducted using the “emmeans” package (Lenth, 2017) to determine the estimated marginal means for the effect of pre-treatment or nectar extraction method between growth type or flower morphology categories, respectively. Coefficients of variation (CV) of nectar traits were calculated to determine the consistency among pre-treatments and nectar extraction methods. These values were calculated by dividing the standard deviation of the nectar trait by its mean, grouped either by growth type and pre-treatment method, or by flower morphology and extraction method. We additionally used Blomberg’s *K* to test for a phylogenetic signal in the three response nectar variables (nectar volume, sugar mass and sugar concentration). Blomberg’s *K* quantifies the variance among phylogenetic clades relative to the variance within clades. The phylogenetic framework was derived from the molecular phylogeny of Zanne et al., (2014) and restricted to the plant genera included in our dataset (Table 1). The tree was generated using the R package “pez” (Pearse et al., 2015) and terminal branches pruned to obtain a cladogram.

### 2.7 Step-by-Step guide

We used the knowledge obtained in our study to develop a step-by-step guide for extracting nectar from various flowers across different growth types and flower morphologies https://doi.org/10.5281/zenodo.22876213. Additionally, we provide a 3D-printable design for a mesh sieve that prevents contamination of nectar by flower parts during centrifugation https://doi.org/10.5281/zenodo.22875506.

## 3. Results

Overall, 654 samples were collected from 23 species (Table 2) across 16 plant families. Of those, 174 samples were from nine woody species amongst eight families, while 480 samples were from 14 herbaceous species across nine families (Table 1). Nectar volumes ranged from an average of 0.02 µl/floret in *Veronica chamaedrys* to up to 9.38 µl/floret in *Berberis verruculosa.* Nectar sugar concentration ranged from an average of 7.33 °Bx (Brix: °Bx = 1g Sucrose/100g solution) in *Deutzia crenata* to up to 74 °Bx in *Berberis verruculosa*.

**Table 2:** The overall summarisation of the means and standard deviation of nectar volume (in µl), sugar concentration (°Brix), and sugar mass (in mg) across all the plant species sampled for this study. The measurements include all pre-treatment groups but solely the measurements using centrifugation extraction method due to the lack of sugar concentration measurements in multiple plant species by means of microcapillary extraction method.

| Species | Nectar vol. (µl) | Sugar conc. (°Bx) | Sugar mass (mg) |
|---|---|---|---|
| *Aegonychon purpurocaeruleum* | 1.19 ± 0.49 | 36.67 ± 8.74 | 2.02 ± 1.04 |
| *Aesculus hippocastanum* | 2.74 ± 2.17 | 16.67 ± 4.63 | 2.15 ± 1.89 |
| *Allium ursinum* | 2.58 ± 2.10 | 16.33 ± 4.92 | 3.48 ± 2.71 |
| *Berberis verruculosa* | 9.38 ± 9.22 | 74.00 ± 2.09 | 7.05 ± 7.06 |
| *Bunias orientalis* | 0.03 ± 0.06 | 21.53 ± 6.37 | 0.05 ± 0.12 |
| *Colutea orientalis* | 1.74 ± 1.18 | 20.47 ± 5.04 | 1.32 ± 0.38 |
| *Cornus sanguinea* | 1.30 ± 1.81 | 29.33 ± 4.14 | 1.75 ± 2.82 |
| *Deutzia crenata* | 0.99 ± 0.45 | 7.33 ± 2.70 | 0.52 ± 0.38 |
| *Dianthus carthusianorum* | 1.65 ± 1.82 | 39.00 ± 3.03 | 0.66 ± 0.76 |
| *Geranium pyrenaicum* | 0.16 ± 0.24 | 10.44 ± 1.75 | 0.03 ± 0.04 |
| *Knautia arvensis* | 3.77 ± 3.29 | 31.56 ± 7.05 | 1.37 ± 0.97 |
| *Kolkwitzia amabilis* | 0.37 ± 0.13 | 26.67 ± 7.62 | 0.79 ± 0.28 |
| *Lamium galeobdolon* | 0.89 ± 0.60 | 24.83 ± 5.70 | 0.47 ± 0.41 |
| *Lotus corniculatus* | 2.20 ± 2.23 | 10.88 ± 2.23 | 1.00 ± 0.93 |
| *Onobrychis viciifolia* | 0.33 ± 0.32 | 26.33 ± 16.39 | 0.74 ± 0.74 |
| *Rosa canina* | 0.71 ± 1.60 | 8.36 ± 1.28 | 0.08 ± 0.16 |
| *Salvia pratensis* | 2.09 ± 4.45 | 50.33 ± 12.86 | 4.14 ± 7.58 |
| *Styrax japonicus* | 1.89 ± 1.13 | 50.67 ± 10.55 | 1.98 ± 1.51 |
| *Trifolium pratense* | 5.21 ± 4.69 | 45.67 ± 4.63 | 2.33 ± 2.15 |
| *Veronica chamaedrys* | 0.02 ± 0.02 | 23.73 ± 1.33 | 0.02 ± 0.02 |
| *Vicia sepium* | 3.64 ± 3.46 | 23.00 ± 16.36 | 8.60 ± 13.24 |
| *Vicia villosa* | 1.44 ± 1.49 | 20.50 ± 16.92 | 2.71 ± 3.29 |
| *Weigela florida* | 1.46 ± 0.76 | 30.17 ± 20.18 | 1.37 ± 1.30 |

### 3.1 Effects of pre – treatment and growth type on nectar traits

Pre-treatments of flowers significantly affected the nectar volume (χ^2^ = 40.54; p = <0.001) and sugar mass (χ^2^ = 49.40; p = <0.001; Table S.1, S.2; Fig. 3). Compared with flowers kept overnight in water, bagged flowers had higher nectar volumes (estimate = 0.992, SE = 0.20, p < 0.001) and sugar masses (estimate = 1.039, SE = 0.22, p < 0.001). Similarly, compared with the control treatment, bagged flowers had higher nectar volumes (estimate = 0.797, SE = 0.19, p < 0.001) and sugar masses (estimate = 1.066, SE = 0.21, p < 0.001; Table S.2; Fig. 3a, b). Growth type did not affect nectar volume and sugar mass (Table S.1). However, sugar concentration was significantly different depending on the pre-treatment used for plants with differing growth types (χ^2^ = 36.02; p = <0.001). For herbaceous plants, pollinator exclusion by bagging resulted in higher sugar concentrations as compared to overnight water storage (estimate = 1.039, SE = 0.22, p < 0.001), and both bagging (estimate = 0.253, SE = 0.05, p < 0.001) and overnight water storage method (estimate = 0.144 SE = 0.05, p = 0.02) contained significantly higher sugar concentrations than the control. For woody species, the overnight water storage (estimate = -0.375, SE = 0.07, p < 0.001) and bagging pre-treatment (estimate = 0.415, SE = 0.09, p <0.001) had significantly higher nectar sugar concentrations as compared to the control.

**Figure 3:**
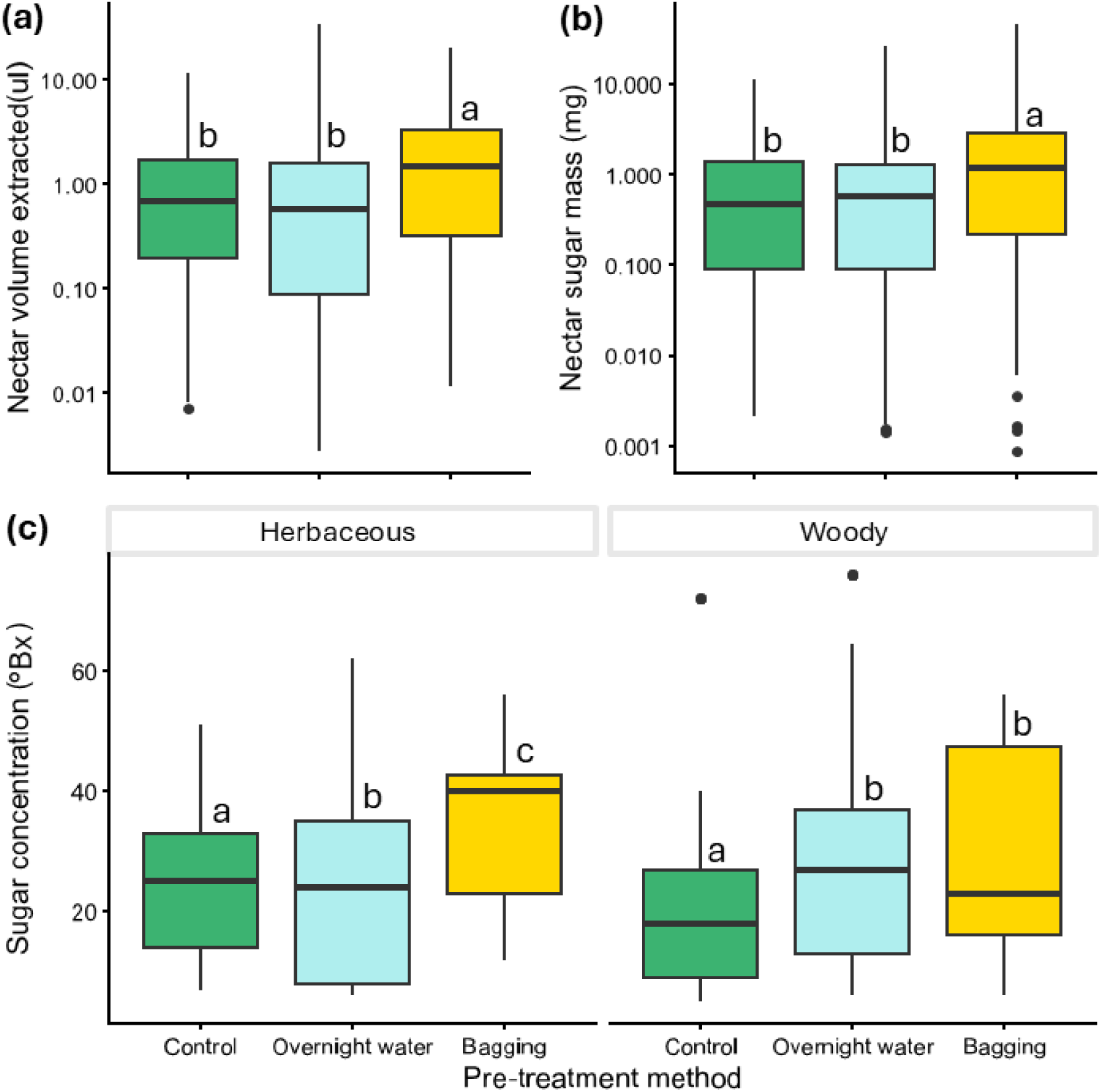
Differences between the three pre-treatments, i.e., pollinator exclusion by bagging (bagging, yellow), overnight water storage (blue), and control (standing crop, green), on nectar measurements. Pre-treatment affects the (a) extractable nectar volume (µl), (b) nectar sugar mass (mg) and (c) nectar concentration (°Bx). Letters on top of the boxplots indicate significant differences between the nectar traits under different pre-treatment conditions prior to extraction.

Although nectar exclusion by bagging resulted in the highest mean nectar volume (Fig. 3a) and nectar sugar mass (Fig. 3b) across growth types as well as the highest mean sugar concentration for herbaceous plants (Fig. 3c), the measurements showed considerable variation, indicating high within-treatment variability, especially for herbaceous species (Table 3). The coefficient of variation (CV) for nectar volume was the highest in woody plants stored overnight in water (2.20) and the lowest for woody plants that were bagged for pollinator exclusion (0.83; Table 3). Overall, for herbaceous plants, nectar volume variation was high (> 1.5) amongst all pre-treatment groups. The variation in sugar concentration for bagged flowers across both growth types (∼ 0.5) was lower than that of plants kept overnight in water (∼ 0.7), while the variation in sugar mass was the highest in bagged herbaceous flowers (2.01) and the lowest for bagged woody plants (0.90; Table 3).

**Table 3:** Coefficients of variation (CV) of the nectar volume (Nec.vol), sugar concentration (Sugar conc.) and sugar mass for different pre-treatment methods grouped by growth types. The pre-treatment groups are control (standing crop), overnight water storage (overnight water), and pollinator exclusion by bagging (bagging).

| Pre-treatment method | Growth type | CV Nec.vol | CV Sugar conc. | CV Sugar mass |
|---|---|---|---|---|
| Control | Herbaceous | 1.69 | 0.51 | 1.50 |
| Control | Woody | 1.11 | 0.84 | 1.45 |
| Overnight water | Herbaceous | 1.90 | 0.66 | 1.26 |
| Overnight water | Woody | 2.20 | 0.70 | 1.96 |
| Bagging | Herbaceous | 1.55 | 0.42 | 2.01 |
| Bagging | Woody | 0.83 | 0.59 | 0.90 |

### 3.2 Effects of extraction method and floral morphology on nectar traits

Across all flower morphology groups, significantly higher nectar volume could be extracted using the centrifugation method as compared to using microcapillaries (χ^2^ = 16.68, p < 0.001; Fig. 4). For plant species with disc-shaped and lip flowers, the nectar extracted using microcapillaries was insufficient for subsequent nectar volume analyses. As at least 5µl of nectar was required for sugar concentration readings using refractometers, sugar could only be quantified for these two groups based on the centrifugation method (Table 2), consequently these two flower morphology groups were removed from the analysis. The nectar sugar concentration was significantly higher for nectar extracted from composite flowers (estimate = 0.371, SE = 0.07, p < 0.001) using the centrifugation method as compared to microcapillaries (Table S.3; Fig. 5a). For stalk disc, papilionaceous, and composite flowers, the sugar mass was significantly higher in nectar extracted using the centrifugation method (Table S.1, S.3; Fig. 5b).

**Figure 4:**
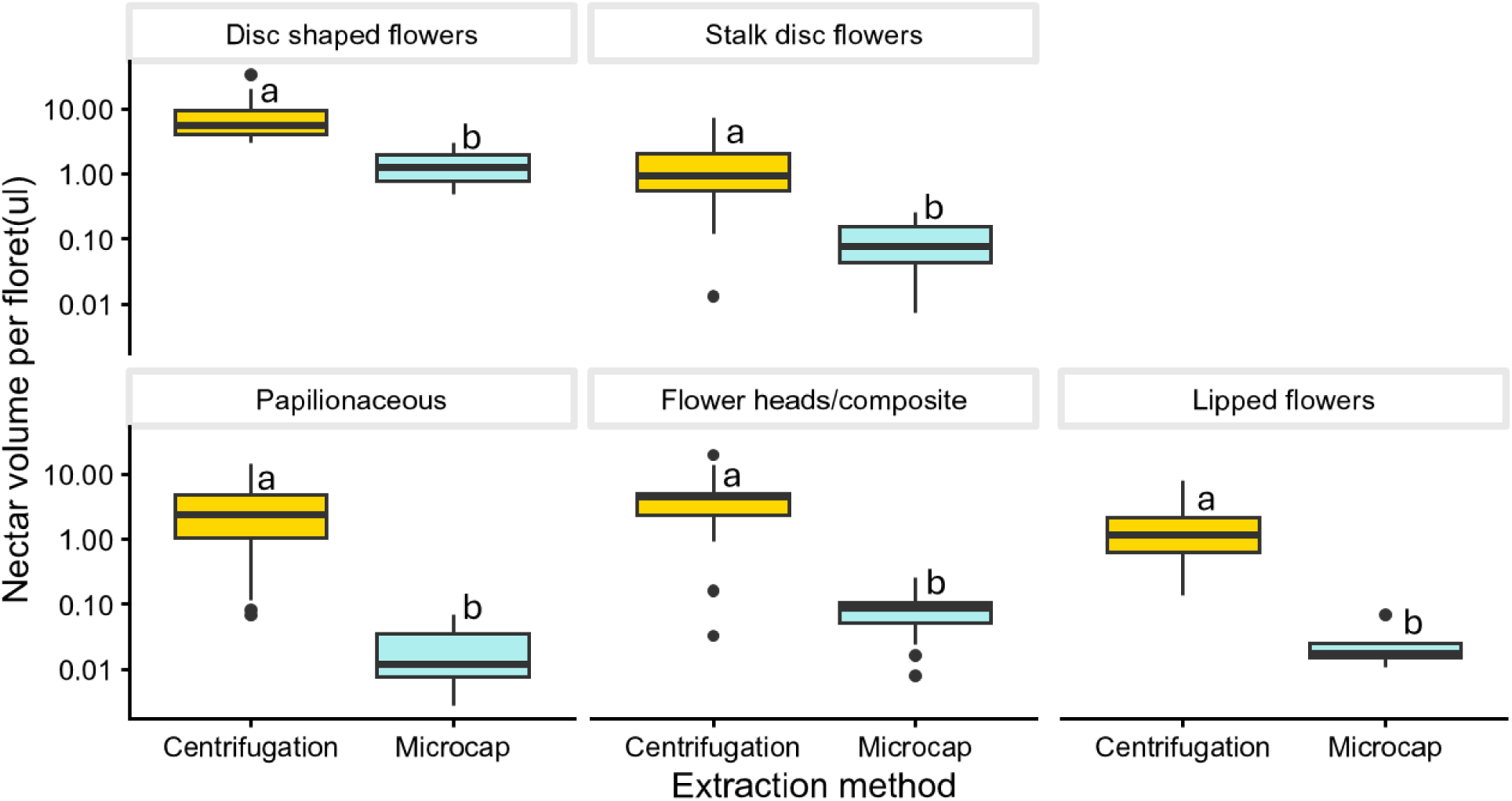
Difference between the nectar volume extracted using two methods, i.e., microcapillary (microcap; depicted in blue) and centrifugation (indicated by yellow), compared amongst various flower morphology categories. Different letters on top of boxplots indicate significant differences in the nectar volume extracted.

**Figure 5:**
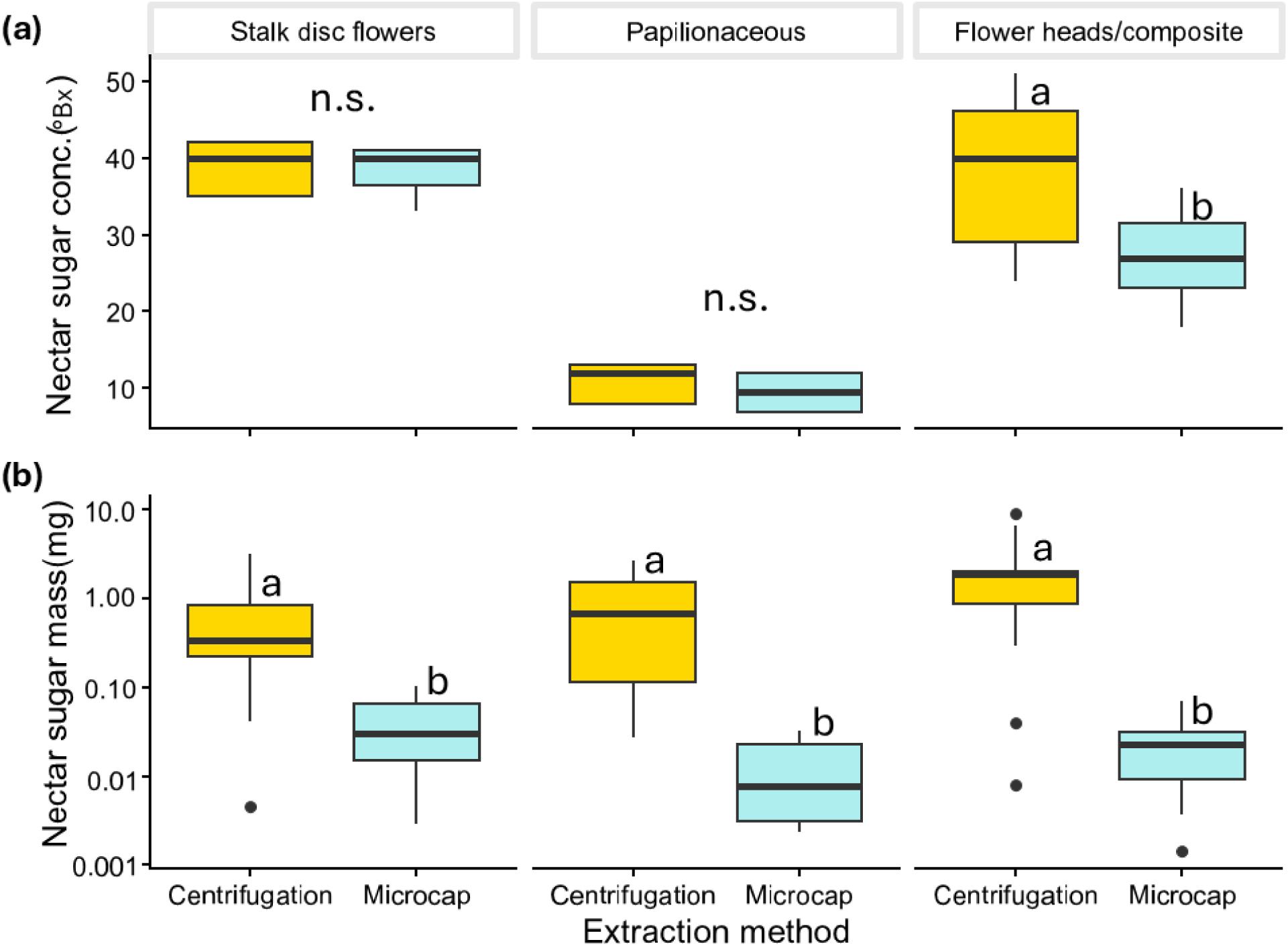
Differences between the sugar concentrations (a) and sugar mass (b) of nectar extracted using two different methods, i.e., microcapillary (microcap; blue) and centrifugation (yellow), compared across various flower morphology categories. Different letters on top of boxplots indicate significant differences in the nectar sugar concentration or mass extracted.

Overall, the variability in nectar volume, calculated as the coefficient of variation, was higher when nectar was extracted using microcapillaries as compared to the centrifugation method for each flower morphology category (Table 4). The variation in sugar concentration was comparable between the extraction methods for composite flowers (0.24 and 0.25 for centrifugation and microcapillary, respectively) but lower for nectar extracted from stalk disc and papilionaceous flowers using microcapillaries as compared to centrifugation (Table 4). The variability in sugar mass of nectar extracted from stalk disc and composite flowers was comparable across extraction methods, but higher for nectar from papilionaceous flowers extracted with the centrifugation method (2.32) as compared to microcapillary extraction (0.93). For sugar concentration and mass measurements, at least 5µl of nectar was required from each species, which could not be obtained using the microcapillary method, for this reason, no CV values could be calculated for species with disc-shaped and lip flowers.

**Table 4:** Coefficients of variation (CV) of the nectar volume, sugar concentration (Sugar conc.) and sugar mass for different nectar extraction methods, i.e., centrifugation and microcapillary, grouped by flower morphology.

| Flower morphology | Extraction method | CV Nectar volume | CV Sugar conc. | CV Sugar mass |
|---|---|---|---|---|
| Disc-shaped | Centrifugation | 1.39 | 0.74 | 1.68 |
| Disc-shaped | Microcap | 2.07 | NA | NA |
| Stalk disc | Centrifugation | 0.95 | 0.38 | 0.87 |
| Stalk disc | Microcap | 0.99 | 0.09 | 0.83 |
| Lip | Centrifugation | 1.85 | 0.52 | 2.40 |
| Lip | Microcap | 2.09 | NA | NA |
| Papilionaceous | Centrifugation | 1.20 | 0.68 | 2.32 |
| Papilionaceous | Microcap | 2.28 | 0.29 | 0.93 |
| Composite | Centrifugation | 0.90 | 0.24 | 0.93 |
| Composite | Microcap | 1.61 | 0.25 | 0.80 |

### 3.3 Effects of phylogeny on nectar traits

Blomberg’s *K* showed a significant phylogenetic signal only for sugar mass (*K* = 2.547, p = 0.002) but not for nectar volume (*K* = 0.623, p = 0.219) or sugar concentration (*K* = 0.518, p = 0.283).

## 4. Discussion

### 4.1 Pre Treatment

The tested pre-treatments - bagging, overnight water storage and control (standing crop) - affected the quantity and quality of extracted nectar. Flowers that were bagged had the highest extractable nectar volume and sugar mass, indicating that bagging successfully prevented nectar consumption by nectivorous animals and resulted in higher nectar accumulation and sampling availability. This is in line with and supports the effectiveness of previous recommendations and common practice (Biella et al., 2022; Corbet, 2003; Jachuła et al., 2021; Venjakob et al., 2022).

The sugar concentrations in nectar showed a similar pattern for both the bagged and control treatments. In general, bagged flowers had higher sugar concentrations than control flowers. Interestingly, for watered flowers, effects on sugar concentrations differed between growth types: in herbaceous plants, sugar concentration was lowest in watered flowers, while in woody plants, sugar concentration of watered flowers was comparable to that of bagged ones (see Fig. 3b, Table S.1, S.2). This might be explained by a lack of additional sugar reserves in herbaceous flowers, whereas woody plants likely have a higher capacity to retain sugar in woody tissues, enabling them to replenish nectar sugar (Hoch, 2005).

### 4.2 Extraction Method

Nectar volume and nectar sugar mass were both higher when a centrifuge was used, though the difference between centrifuging and microcapillaries varied with flower shape. This finding demonstrates, as expected, that centrifuging can extract more nectar (and thus more nectar sugar) from a broad selection of flower shapes compared with the capillary method. Moreover, it was not possible to extract nectar with capillaries for several flowers, as even a 0.5 µl capillary was too thick for their corolla, or the nectar was provided in a too shallow pit, preventing uptake by capillaries. We also could not measure sugar concentrations for flowers with low nectar volumes. Interestingly, even flowers with deeper corollas (e.g., Papilionaceous species such as *Vicia sepium*, *Vicia villosa,* or *Lotus corniculatus*) and tube-shaped nectaries or nectar-reservoir compartments showed marked differences in nectar volumes extracted with different methods. In contrast, we found comparatively less variation among extraction types for shallower flowers (disc-shaped flowers, e.g., *Cornus sanguinea* or *Allium ursinum*), which could reflect either low measurements overall, a result of sampling with more pooled flowers per measurement, or experimenter bias. To conclude, all flower shapes tested in our study yielded greater extraction volumes and sugar mass with the centrifuge method, which may therefore provide a more precise estimate for nectar volume measurements.

Notably, and contrary to expectations, nectar sugar concentration was also affected by the sampling method. However, this difference might be explained primarily by a comparatively lower nectar concentration in capillary-extracted flowers within the “Flower heads/composite” category, most likely due to dilution caused by plant damage during capillary use. These flowers (e.g., *Trifolium pratense* & *Knautia arvensis*) typically have tiny florets and are easily damaged with the thin capillaries. Except for floral shape (simplified Kugler category), there were no significant differences between extraction methods, suggesting that nectar quality was hardly affected by the extraction method.

Overall, we consider the centrifugation method the best approach for measuring nectar volume, sugar mass, and sugar concentration, contradicting the conclusions of Pioltelli et al., (2024), who reported lower volumes with centrifugation. This discrepancy might arise from the flowering species used, as three bigger ornamental flowers were measured and not further specified methodological details.

Interestingly, we did not find a phylogenetic signal for either nectar volume or sugar concentration, but for sugar mass. Possibly both nectar sugar concentration and nectar volume are stronger linked to pollinators and morphological constraints compared to the sugar mass (Parachnowitsch et al., 2019). As our sample size didn’t allow for further complexity, we refrained from making any inferences on the effect of the extraction method on nectar sugar mass from being drawn in this study.

### 4.3 Limitations

A potential bias driving differences between our pre-treatment method watering was the overnight location, as plants were kept under lab conditions, resulting in altered climatic conditions (temperature, humidity) that differed from those outside. Additionally, bagging not only leads to an accumulation of sugar in flowers but also increases humidity inside bags (Wyatt et al., 1992) which has the potential to alter results. However, we did not observe a lower nectar sugar concentration in bagged flowers, indicating that this factor is negligible in our study.

During extraction, additional bias could have been induced by plant tissue damage. However, nectar sugar concentration was higher, not lower (as would be expected following tissue damage) in the centrifuged flowers, indicating less or no contamination of nectar by other tissue fluids, as compared to the capillary method. Notably, for some plant species, we could not extract sufficient nectar for refractometer analyses, particularly from capillary samples, resulting in insufficient sample sizes for some comparisons. Our study focused on the effects of different pre-treatments and extraction methods on nectar volume, sugar mass and sugar concentration. The applied methods and suggested protocols (see below) can be further adapted and expanded in future studies, e.g., by including additional plant species or by analysing the chemical composition of nectar. While we included as many local plant species and morphologies as possible, we need to point out that there is a plethora of plants for which nectar extraction is simply not possible without manipulation/mobilisation of the nectar through additional water or a solvent (Ethanol), e.g., common ivy (*Hedera helix*), which has an almost crystallised nectar structure.

Below, we provide best practice guidelines and a step-by-step guide based on our findings:

#### Pre-treatments

We suggest selecting pre-treatments based on the question at hand. For example, morning collection of undisturbed plants (control) was quite successful and is probably the most appropriate approach for assessing, e.g., daily nectar availability for foragers, location-specific effects or gradient effects. Bagging has a distinct bias towards higher volumes and should be used to assess the maximum amount of nectar a flower can produce. It can also support subsequent chemical analyses that require larger nectar volumes. In our samples, sugar concentrations were higher in bagged flowers than in the control (standing crop), indicating that bagging not only influences volume but also composition, an important bias to consider. In general, we advise against watering flowers overnight, especially under uncontrolled overnight conditions (e.g., not in a temperature and humidity-controlled room) as we couldn’t find a consistent positive improvement with this method.

#### Extraction

Based on our results, we recommend the centrifuge method wherever possible. Not only does it yield the highest nectar amounts, but it also enables easy and fast nectar extraction, especially using 3D printed sieves, which prevent debris from contaminating the collected nectar. When field sites are located far away from a laboratory, it is also possible to bring centrifuges, even for 50ml Falcon tubes, and portable batteries to the field locations and run them, e.g. in cars. There may, however, be good use cases for capillary-collected nectar, e.g., if flowers cannot be harvested in the field.

## 5. Conclusion

Our study highlights the importance of the treatment of flowering plants prior to nectar extraction to avoid biases and introduces a standardised approach to centrifuge-based nectar extraction using additional 3D-printed universal sieves. This approach resulted in not only higher nectar volumes but also enabled, standardised sampling of nectar from a broad range of plant species. This method could enable or simplify future in-depth analysis of nectar compositions, thus supporting future studies on the role of nectar in plant-pollinator interactions and floral biology.

## Authors contributions

V.B., C.A.N, P.P., K.K., G.T.K., S.D.L. conceived the ideas and designed methodology; A.D.Ö and L.X. collected the data; V.B., P.P., K.K., G.T.K., F.A.R. analysed the data; V.B. and K.K. led the writing of the manuscript. All authors contributed critically to the drafts and gave final approval for publication.

## Statement of Inclusion

This study was carried out by both local and international scientists with varying levels of experience, in a collaborative effort.

## Data availability statement

Data and the R script used for this study are available under the following link to Zenodo: https://doi.org/10.5281/zenodo.22876213. The models for the 3D-Prints of the sieves used in this study are similarly available under the following link to Zenodo: https://doi.org/10.5281/zenodo.22875506. A step by step guide for the adaptive centrifuge extraction of floral nectar based on the results of this study are also available under the following Zenodo link: https://doi.org/10.5281/zenodo.24201322.

## Acknowledgment

We want to acknowledge the contribution of both A. S. Höwener and A. E. Neumann to the development of parts of the described centrifuge method. The work of K. Kuba was supported by the German Federal Environmental Foundation [20022/14].

