## Supplementary material for "Quantifying floral rewards: Comparative evaluation of nectar extraction techniques"

**Table S.1**: Results from generalised linear mixed-effect models (GLMMs) to determine the interactive effects of pre-treatment and growth type, or nectar extraction technique and floral morphology on nectar trait measurements. The explanatory and response variables, AIC (Akaike information criterion), conditional and marginal R² (obtained via the “MuMIn” package), chi-square (χ²), degrees of freedom as (df), and p-values (p), are described, with significant p-values marked in bold. For all models, the species, temperature and the weather at the time of sample collection were included as random variables.

| **Response variables** | **N** | **Marginal R²** | **Conditional R²** | **Model Predictors** | **χ^2^** | **df** | **p** |
| --- | --- | --- | --- | --- | --- | --- | --- |
| Nectar volume per floret | 436 | 0.037 | 0.816 | Pre-treatment | 40.54 | 2 | **<0.001** |
|  |  |  |  | Growth type | 0.90 | 1 | 0.343 |
|  |  |  |  | Pre-treatment * Growth type | 1.21 | 2 | 0.545 |
| Nectar sugar concentration | 424 | 0.052 | 0.801 | Pre-treatment | 59.04 | 2 | <0.001 |
|  |  |  |  | Growth type | 0.02 | 1 | 0.880 |
|  |  |  |  | Pre-treatment * Growth type | 36.02 | 2 | **<0.001** |
| Nectar sugar mass | 424 | 0.065 | 0.691 | Pre-treatment | 49.40 | 2 | **<0.001** |
|  |  |  |  | Growth type | 0.46 | 1 | 0.499 |
|  |  |  |  | Pre-treatment * Growth type | 2.59 | 2 | 0.275 |
| Nectar volume per floret | 179 | 0.631 | 0.712 | Extraction method | 6.47 | 1 | 0.011 |
|  |  |  |  | Flower shape | 24.02 | 4 | <0.001 |
|  |  |  |  | Extraction method * Flower shape | 16.68 | 4 | **<0.001** |
| Nectar sugar concentration | 101 | 0.708 | 0.895 | Extraction method | 0.04 | 1 | 0.839 |
|  |  |  |  | Flower shape | 19.31 | 2 | <0.001 |
|  |  |  |  | Extraction method * Flower shape | 10.55 | 2 | **0.005** |
| Nectar sugar mass | 101 | 0.591 | 0.657 | Extraction method | 30.63 | 1 | <0.001 |
|  |  |  |  | Flower shape | 2.65 | 2 | 0.266 |
|  |  |  |  | Extraction method * Flower shape | 8.45 | 2 | **0.015** |

**Table S.2**: Results from the pairwise comparisons of estimated marginal mean with Tukey’s adjustment to determine the differences in nectar traits among the pre-treatment groups – pollinator exclusion by bagging (bagging), standing crop (control) and overnight water storage (overnight water). The table summarizes chi-square values (χ²), standard error (SE), degrees of freedom as (df), and Tukey-adjusted p-values (p), with significant p-values marked in bold.

| **Response variable** | **Pre-treatment method** | **χ²** | **SE** | **df** | **t-ratio** | **p-value** |
| --- | --- | --- | --- | --- | --- | --- |
| Nectar vol | Bagging - Control | 0.797 | 0.19 | 420 | 4.17 | **<0.001** |
|  | Bagging - Water storage | 0.992 | 0.20 | 421 | 5.02 | **<0.001** |
|  | Control - Water storage | 0.194 | 0.16 | 405 | 1.22 | 0.449 |
| Sugar conc. - Herbaceous | Bagging - Control | 0.253 | 0.05 | 398 | 4.98 | **<0.001** |
|  | Bagging - Water storage | 0.397 | 0.05 | 385 | 7.40 | **<0.001** |
|  | Control - Water storage | 0.144 | 0.05 | 401 | 2.72 | **0.019** |
| Sugar conc. - Woody | Bagging - Control | 0.415 | 0.09 | 243 | 4.60 | **<0.001** |
|  | Bagging - Water storage | 0.040 | 0.10 | 195 | 0.42 | 0.908 |
|  | Control - Water storage | -0.375 | 0.07 | 399 | -5.54 | **<0.001** |
| Sugar mass (mg) | Bagging - Control | 1.066 | 0.21 | 116 | 4.98 | **<0.001** |
|  | Bagging - Water storage | 1.039 | 0.22 | 100 | 4.65 | **<0.001** |
|  | Control - Water storage | -0.027 | 0.17 | 395 | -0.16 | 0.987 |

**Table S.3**: Results from the pairwise comparisons of estimated marginal mean for each extraction method, grouped by flower morphology, with Tukey’s adjustment. The table summarizes chi-square values (χ²), standard error (SE), degrees of freedom as (df), and Tukey-adjusted p-values (p), with significant p-values marked in bold.

| **Response variable** | **Flower shape** | **Extraction method** | **χ²** | **SE** | **df** | **t-ratio** | **p-value** |
| --- | --- | --- | --- | --- | --- | --- | --- |
| Nectar vol | Disc/bowl shaped | Centrifugation - Microcapillary | 2.180 | 0.87 | 166 | 2.51 | **0.013** |
|  | Stalk disk |  | 2.480 | 0.39 | 163 | 6.41 | **<0.001** |
|  | Lipped |  | 3.990 | 0.63 | 32 | 6.35 | **<0.001** |
|  | Papilionaceous |  | 4.570 | 0.47 | 167 | 9.74 | **<0.001** |
|  | Composite |  | 3.880 | 0.34 | 164 | 11.29 | **<0.001** |
| Sugar conc. (Brix) | Disc/bowl shaped | Centrifugation - Microcapillary | NA | NA | NA | NA | NA |
|  | Stalk disk |  | 0.013 | 0.07 | 91 | 0.20 | 0.840 |
|  | Lipped |  | NA | NA | NA | NA | NA |
|  | Papilionaceous |  | 0.089 | 0.09 | 93 | 0.95 | 0.344 |
|  | Composite |  | 0.371 | 0.07 | 92 | 4.52 | **<0.001** |
| Sugar mass (mg) | Disc/bowl shaped | Centrifugation - Microcapillary | NA | NA | NA | NA | NA |
|  | Stalk disk |  | 2.490 | 0.45 | 92 | 5.54 | **<0.001** |
|  | Lipped |  | NA | NA | NA | NA | NA |
|  | Papilionaceous |  | 4.130 | 0.65 | 92 | 6.36 | **<0.001** |
|  | Composite |  | 4.200 | 0.49 | 82 | 8.58 | **<0.001** |
